# Kappa Opioid Receptors Negatively Regulate Real Time Spontaneous Dopamine Signals by Reducing Release and Increasing Uptake

**DOI:** 10.1101/2024.02.05.578840

**Authors:** Conner W Wallace, Katherine M Holleran, Clare Y Slinkard, Samuel W Centanni, Sara R Jones

## Abstract

The role of the dynorphin/kappa opioid receptor (KOR) system in dopamine (DA) regulation has been extensively investigated. KOR activation reduces extracellular DA concentrations and increases DA transporter (DAT) activity and trafficking to the membrane. To explore KOR influences on real-time DA fluctuations, we used the photosensor dLight1.2 with fiber photometry in the nucleus accumbens (NAc) core of freely moving male and female C57BL/6 mice. First, we established that the rise and fall of spontaneous DA signals were due to DA release and reuptake, respectively. Then mice were systemically administered the KOR agonist U50,488H (U50), with or without pretreatment with the KOR antagonist aticaprant (ATIC). U50 reduced both the amplitude and width of spontaneous signals in males, but only reduced width in females. Further, the slope of the correlation between amplitude and width was increased in both sexes, suggesting that DA uptake rates were increased. U50 also reduced the frequency of signals in both males and females. All effects of KOR activation were stronger in males. Overall, KORs exerted significant inhibitory control over spontaneous DA signaling, acting through at least three mechanisms - inhibiting DA release, promoting DAT-mediated uptake, and reducing the frequency of signals.

## 1. Introduction

The kappa opioid receptor (KOR) system exerts a strong inhibitory influence over dopamine (DA) neurotransmission. A primary outcome of KOR activation in the nucleus accumbens (NAc) is a reduction in extracellular DA concentrations, as shown using *in vivo* microdialysis [1, 2]. DA levels are maintained by a balance between release and uptake, and KORs regulate both processes. Using *ex vivo* fast scan cyclic voltammetry (FSCV), our lab has shown that KOR agonists dose-dependently reduce electrically stimulated DA release in NAc brain slices [3–5], and that this effect is heightened by stress or chronic drug or alcohol exposure [6, 7]. KOR-mediated reductions in DA release are well-documented [8–10], but KOR effects on DA uptake are less consistent. Several reports showed that KOR activation increased DAT trafficking and uptake rates in cultured cells and tissue homogenates [11, 12], and in the striatum of anesthetized animals [13], but no uptake changes were measured in NAc core slices [4, 5]. Extensive work from the Shippenberg lab reported that acute administration of KOR agonists had no discernable effects on DA uptake, but repeated administration significantly reduced the maximal rate of uptake [14, 15]. Thus, ample evidence demonstrates that KORs regulate DA, but its mechanisms of action are not well-defined. Methodological differences between reports may be making the elucidation of precisely how KORs affect DA terminals more difficult.

KOR activation has also been shown to inhibit the firing of VTA DA neurons projecting to the NAc [16] and prefrontal cortex [17]. Binding of the KOR by its ligands activates G-protein-coupled inwardly rectifying potassium channels in cultured cells [18] and in DA neurons [19], resulting in hyperpolarization that directly inhibits firing. Alterations in firing could affect DA signaling in the NAc, such as changing the magnitude of individual release events or basal extracellular concentrations [20]. The influence of KORs over DA is a complex interplay of altered cell body firing, terminal release, and uptake changes. The combined effects of these mechanisms are not well understood. Therefore, we measured how KORs regulate spontaneous DA neurotransmission in intact animals on a rapid time scale to obtain a more holistic assessment.

Assessment of spontaneous DA signal kinetics is relatively new. Since the advent of the dLight photosensor [21], intense focus has been given to how DA in the NAc responds to salient cues and controls specific mechanics of learning [22–25]. Decades of work with established techniques (microdialysis, voltammetry, electrophysiology) have elucidated the kinetics of electrically- and cue-evoked DA signaling and firing patterns of DA neurons *in vivo* in response to acute, experimenter-administered or repeated self-administration of alcohol and other drugs of abuse [26–30]. Investigators recently have built on this work by assessing DA responses using dLight during alcohol self-administration [31], and in response to stimulants [32–35] and opioids [33, 36, 37]. While FSCV studies in slices [38, 39] and *in vivo* [30, 40] have noted the presence of spontaneous DA signals, it has never been possible to assess these events in awake and behaving animals on such a rapid time scale as with dLight. It was recently shown that the width of spontaneous dLight signals differs across regions of the striatum [41] and that the amplitude is increased after repeated naloxone-precipitated withdrawal [36]. To assess the kinetics of spontaneous signals and determine how KOR activation might alter them, we evaluated these variables (amplitude and width) in a novel analysis method. Our lab [42] and others [43, 44] have shown with FSCV in brain slices that DA release and uptake are highly correlated. Therefore, we decided to pharmacologically manipulate DA release and uptake to model how this relationship might be explained using the amplitude and width of dLight signals, including the correlation between them.

In this study, using fiber photometry paired with the fluorescent DA photosensor dLight in awake and freely behaving mice, modulation of spontaneous DA signals using KOR pharmacology was examined for the first time. We hypothesized that KOR activation by U50,488H (U50) would reduce release, which would be reflected in lower peak amplitudes of dLight signals, with little or no effects on uptake, measured primarily by signal width, and that these effects would be blocked with the KOR antagonist aticaprant (ATIC), which would have no effects on its own. Additionally, because female rodents have been shown to be less sensitive to KOR agonist-induced depressive behavior and inhibition of DA release [45, 46], we predicted that U50 would reduce signal amplitude less robustly in females. Indeed, the results showed that U50 reduced signal amplitude in males but not females. However, contrary to expectations, width was decreased in both sexes, with the magnitude of decrease in width being greater than the reduction in amplitude in males. When signal amplitude and width were correlated, U50 increased the slope in both sexes, confirming increased uptake rates. U50 also reduced signal frequency in both sexes. All U50 effects were reversed by ATIC, but unexpectedly, ATIC alone increased DA signal amplitude in both males and females, suggesting that there is tonic KOR activity in normal, untreated mice. Overall, this investigation found that KORs inhibit DA signaling *in vivo* through multiple mechanisms that differ between sexes, and it presents a novel method by which to identify and assess spontaneous DA signaling events in real time.

## 2. Materials and Methods

### 2.1. Animals

Animal protocols were reviewed and approved by the IACUC. C57BL/6J mice (n

= 7 male, n = 7 female; Jackson Laboratory, Bar Harbor, ME) acclimated to housing facilities for at least one week before surgery. Housing lights were 12h:12h, on at 0600. Water and standard rodent chow were available *ad libitum*. Mice were ∼12 weeks old at the start of behavioral testing.

### 2.2. Stereotactic surgery

Mice were induced and maintained under anesthesia with isoflurane and received 5 mg/kg meloxicam before surgery. Surgeries were performed on a computerized stereotactic frame from Neurostar (Tübingen, Baden-Württemberg, Germany). The dLight virus (700 nL of AAV5-hSyn-dLight 1.2; Addgene, Watertown, MA; Catalog No.111068-AAV5) was injected unilaterally into the NAc core (in mm: AP:+1.1, ML: ± 1.3, DV: +4.6 from Bregma). Afterwards, a fiber optic cannula (Doric Lenses, Québec, Canada) was inserted to a depth of 0.1 mm above the injection site. Adhesive cement (C&B-Metabond, Parkell, Edgewood, NY, Part No.S380) was applied to secure the cannula and close the surgical site followed by Neo-Predef with Tetracaine antibiotic powder (Zoetis, Parsipanny, NJ). Mice were monitored and administered meloxicam daily for 2 more days as needed for pain.

### 2.3. dLight fiber photometry

Photometry equipment was from Fiber Photometry Systems at Tucker Davis Technologies (TDT) (RZ5P, Synapse, Alachua, FL) and Doric Lenses (Québec, Canada). Two excitation wavelengths, 405 nm (isosbestic control signal) and 465 nm (dLight1.2-dependent signal), were emitted from LEDs (CLED405, CLED465, Doric Lenses) and controlled by a multi-channel programmable LED driver (LEDD_4, Doric Lenses). These signals were channeled into 0.57NA, 400μm core, pre-bleached, low-autofluorescence patch cords using a Doric 4-Port Fluorescence Mini Cube (FMC4, Doric Lenses). Fluorescence signals were detected using a Visible Femtowatt Photoreceiver (Newport Corporation, Irvine, CA; Model 2151). Synapse software was used to modulate excitation lights (405 nm: 210 Hz, 465 nm: 330 Hz) and to measure demodulated and low-pass filtered (6 Hz) fluorescence signals that were transduced in real time via the RZ5P processor. Four weeks postoperatively, an initial test was performed to set light intensity levels for each animal. The 405 nm light was maintained at 15 μW for all animals. The intensity of the 465 nm light was gradually increased from 15 to 30 μW until clear signals were seen. Mice without discernible signals were excluded.

### 2.4. Pharmacology

Mice were acclimated inside operant boxes (Med Associates, Inc., Fairfax, VT, Part No. ENV-307A) for 1 hour/day for 5 days. Mice underwent two sessions weekly, preceded by saline or drug injections. Drugs included U50 (5 mg/kg, i.p.), ATIC (10 mg/kg, s.c.), saline (i.p.), and ATIC vehicle control (s.c.). Saline and U50 were administered 30 minutes before recordings, and mice were tethered immediately after these injections for acclimation in the boxes. ATIC and its vehicle were given 1 hour before the session, and mice were returned to the home cage for 30 minutes prior to saline or U50 injection. The U50 alone session was performed before sessions with ATIC to prevent antagonist-mediated alterations to KOR sensitivity, which have been shown with other KOR antagonists [47].

A subset of mice (male: n = 3; female: n = 3) was randomly chosen to complete a series of proof-of-concept experiments for kinetic analyses of spontaneous DA signals after KOR pharmacology trials. Sessions were preceded either by saline (15 minutes), cocaine (immediately, 10 mg/kg), raclopride (15 minutes, 0.3 mg/kg), or SCH23390 (30 minutes, 1 mg/kg) (i.p.). The order was counterbalanced, but SCH23390 was given last to prevent signal attenuation. Mice completed four total 10-minute home cage recordings between several days of washout.

2.5. Drugs

Aticaprant (MedChemExpress, Monmouth Junction, NJ; Product: HY-101718, CAS: 1174130-61-0) was dissolved in 5% DMSO / 5% Cremophor / 90% saline at 2 mg/mL. U50,488H (RTI Log No.: 13692-54B) and cocaine ((-)-cocaine HCl, RTI Log No.: 14201-134), generously provided by the NIH, were dissolved in saline at 1 mg/mL and 2 mg/mL, respectively. Raclopride (S(-)-raclopride (+)-tartrate salt, CAS: 98185-20- 7, Sigma Aldrich, Product No.: R121) was dissolved in saline at 0.12 mg/mL. SCH23390-HCl (CAS: 125941-87-9) (Tocris, Product No.: 0925) was dissolved in saline at 1 mg/mL.

### 2.6. dLight fiber photometry analysis

Raw data from Synapse was extracted into a format readable by MATLAB (v. R2023a) using code from TDT. Raw 465 nm fluorescence was normalized to the 405 nm signal and transformed into ΔF/F values. This ΔF/F trace was input into AXON pCLAMP software (Clampfit v. 11), and the baseline was manually adjusted for drift. Control trials were assessed to determine a ΔF/F cutoff, specific for each animal, by which to assess drug effects. A cutoff was chosen that would exclude the baseline ‘noise’ (where positive and negative deflections surrounding ΔF/F = 0 have similar amplitude). The amplitude, “width,” and peak-to-peak frequency of these signals was subsequently assessed, with “width” defined as the duration in seconds from when the signal exceeded to when it fell back below the cutoff.

To compare sex effects and account for differences in viral expression *between* animals during saline control trials, code developed in MATLAB by Martianova et al. [48] and Zhang et al. [49] was used. This automatically transforms ΔF/F values to remove drift followed by a z-score normalization using an adaptive iteratively reweighted Penalized Least Squares (airPLS) algorithm. A standard z-score threshold of z=2.6 (p<0.01) was set for spontaneous signals.

### 2.7. Statistical analysis

Data were analyzed in each sex separately using GraphPad Prism (v. 9.5.1.). Amplitude and width values were normalized within each animal to saline or vehicle trials. One-way analysis of variance (ANOVA) or two-tailed t-tests compared effects of KOR pharmacology on amplitude, width, and frequency. Amplitude and width were correlated, and the slopes of the lines were compared using simple linear regression. Slopes for each drug within each animal were used to compare between drug conditions using paired t-tests. Šídák’s multiple comparisons test was used to compare saline, U50, and ATIC + U50 trials, and unpaired t-tests were used for ATIC and vehicle trials. In all cases where data are directly compared, values for the vehicle are presented before the drug (i.e. vehicle vs. drug; p<0.05).

## 3. Results

### 3.1. Effects of DA-modulating drugs on spontaneous dLight signals

Figure 1 shows that spontaneous dLight signals in the NAc are due to activation of the dLight receptor and that the peak magnitude and rate of return to baseline can be primarily attributed to DA release and uptake, respectively. Representative dLight traces are shown from an animal that received saline or the DA D1 receptor (D1R) antagonist SCH23390 (1 mg/kg) (Figure 1A). A paired t-test showed signal frequency was significantly reduced by SCH23390 (0.29 ± 0.0046 vs. 0.00063 ± 0.00063 Hz; p<0.0001) (Figure 1B). The DA-responsive moiety in dLight is a modified D1R, and signals were blocked by SCH23390, confirming that spontaneous signals resulted from dLight activation.

**Figure 1:**
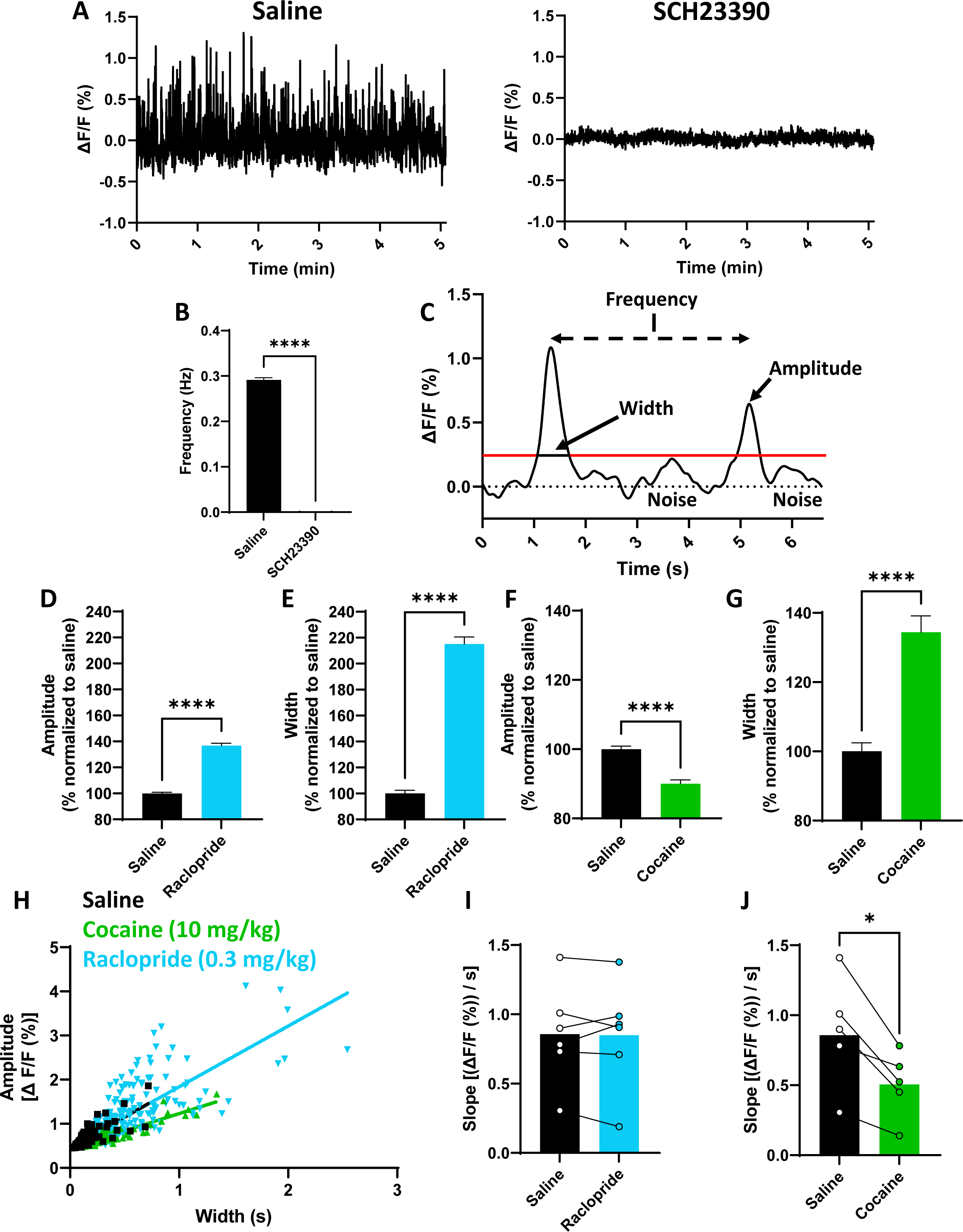
Effects of DA-modulating drugs on spontaneous dLight signals. Two 5-minute traces from one animal show that blocking dLight with a D1R antagonist (SCH23390) prevents spontaneous signals, confirming changes in dLight are from DA **(A).** When repeated in multiple mice (n=3 males, n=3 females), SCH23390 fully blocks signals, as seen by a 99% reduction in frequency (Hz) **(B).** An example trace shows two signals and their ΔF/F cutoff with the measured dependent variables in the study: amplitude, width, and frequency **(C).** Raclopride increased the amplitude **(D)** and width **(E)** of spontaneous signals, whereas cocaine reduced the amplitude **(F)** but alternately increased width **(G)** (all normalized to saline control trials). Amplitude and width were correlated for each trial within each animal, with an example from one animal shown in **(H).** Raclopride did not alter the slope (expressed as [(ΔF/F (%)) / s]) of this correlation **(I)**, but cocaine reduced the slope **(J)**. This suggests raclopride increased dopamine release but cocaine slowed uptake, confirming two known effects of these drugs. Thus, fiber photometry and dLight can be used to assess kinetics of spontaneous DA signals. (*, p&lt;0.05; ****, p&lt;0.0001; significant difference between saline and drugs).

Figure 1C shows a representative trace of two spontaneous signals illustrating amplitude, width, and frequency. The ΔF/F cutoff (shown in red) differentiates signal from noise. The DA D2-like receptor (D2R) antagonist raclopride blocks presynaptic DA autoreceptors, releasing terminals from inhibition and increasing exocytotic release [40]. This should result in an overall larger signal, with a correspondingly higher peak and larger width. As expected, raclopride (0.3 mg/kg) increased signal amplitude (100.0 ± 0.9% vs. 136.8 ± 1.8%; p<0.0001) (Figure 1D) and width (100.0 ± 2.5% vs. 215.2 ± 5.4%; p<0.0001) (Figure 1E). Conversely, cocaine, which is an uptake inhibitor, reduces clearance and increases extracellular DA concentrations [50], but at higher doses it reduces exocytotic release through autoreceptor activation [51, 52]. Cocaine was therefore expected to produce shorter, wider DA signals. Indeed, cocaine (10 mg/kg) reduced signal amplitude (100.0 ± 0.9% vs. 90.1 ± 1.1%; p<0.001) (Figure 1F) and increased width (100.0 ± 2.5% vs. 134.4 ± 4.7%; p<0.001) (Figure 1G).

In NAc-containing brain slices, DA release and uptake were shown to be strongly correlated [42]. We aimed to explore this relationship using amplitude and width to determine if a similar relationship was present and similarly modulated by compounds that specifically alter release or uptake. If a drug primarily affects release, width should scale with amplitude, resulting in no change in the slope of a line depicting the correlation between amplitude and width. However, if a drug is applied which promotes or inhibits uptake, the slope will increase or decrease, respectively. A representative example within one animal is shown in Figure 1H. There was no effect of raclopride on slope (0.856 ± 0.148 vs. 0.849 ± 0.159 %ΔF/F / s; p=0.8777) (Figure 1I), but cocaine reduced the slope (0.856 ± 0.148 vs. 0.506 ± 0.107 %ΔF/F / s; p=0.0163) (Figure 1J). Together, these data indicate a primary effect of raclopride on release and of cocaine on uptake, as was predicted, and highlight the utility of slope as a proxy for DA uptake.

### 3.2. Sex differences in release and uptake

Next, we examined potential sex differences in spontaneous dLight signal kinetics during saline trials. When normalized to the male group, signals from females had a larger amplitude (males: 100.0 ± 0.5%; females: 106.5 ± 0.6%; p<0.0001) (**Figure S1A**), but no difference in width (males: 100.0 ± 1.3%; females: 102.8 ± 1.2%; p=0.1061) (**Figure S1B**). In **Figure S1C**, amplitude values are plotted against width with slope shown as a regression line for each sex. Females had a significantly greater slope (males: 9.671 z-score/s; females: 12.75 z-score/s; p<0.0001) (**Figure S1D**), suggesting that females have faster DA uptake.

### 3.3. KOR activation reduces DA signals through both release and uptake mechanisms

To probe KOR system regulation of DA neurotransmission, the KOR agonist U50,488H (U50, 5 mg/kg, i.p.) was administered with or without pretreatment with the KOR antagonist aticaprant (ATIC, 10 mg/kg, s.c.) prior to dLight recordings. In males, one-way ANOVA revealed a treatment effect on signal amplitude (F(2,6218) = 19.68; p<0.0001) (Figure 2A). Post-hoc tests showed that U50 reduced signal amplitude (100.0 ± 0.9% vs. 88.6 ± 1.4%; p<0.0001), and this reduction was blocked by ATIC (94.7 ± 1.2%; p<0.01). There was also a treatment effect on signal width (F(2,6218) = 32.14; p<0.0001) with a reduction by U50 (100 ± 1.3% vs. 78.3 ± 2.0 %; p<0.0001) that was blocked by ATIC (96.7 ± 1.7%; p<0.0001) (Figure 2B). As shown by a repeated-measures one-way ANOVA, this resulted in a treatment effect on the slope between amplitude and width (F(1.128,4.511) = 6.857; p = 0.0506) (Figure 2C). Post-hoc analysis revealed that U50 increased the slope compared to saline (0.7380 ± 0.1159 vs. 0.9449 ± 0.1082 %ΔF/F / s; p = 0.0199) which was blocked by ATIC (0.6312 ± 0.1137 %ΔF/F / s; p = 0.0390). Thus, KOR activation in males reduces the amplitude of spontaneous signals and increases DA uptake rates.

**Figure 2:**
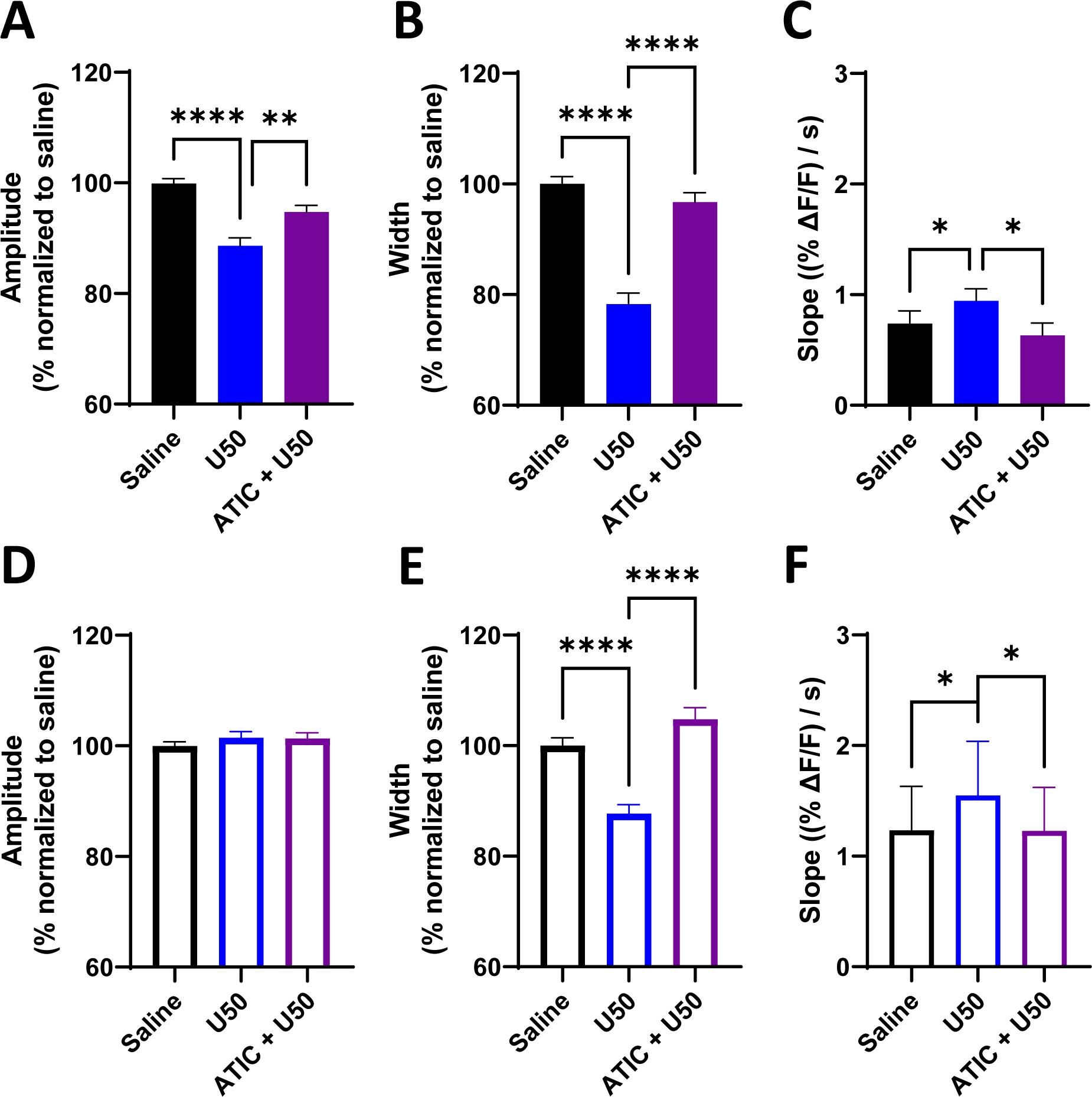
KOR activation reduces DA signals through both release and uptake mechanisms. Using the same analysis principles established in Figure 1, KOR modulating drugs were administered to assess pharmacological effects on dLight signals. In males, KOR activation with U50 reduced the amplitude (A) and width (B) of spontaneous signals (normalized to saline control trials) but increased the slope [(ΔF/F (%)) / s] of the correlation between these two variables (C). This indicates KOR-mediated reductions in DA release with faster DA uptake. In females, U50 did not reduce signal amplitude (D), but it did reduce signal width (E) and increased the slope (F). This suggests KOR activation in females hastens the rate of DA uptake, similarly to males, but it does not reduce DA release. All significant effects seen due to U50 were blocked by pretreatment with the KOR antagonist ATIC. (*, p&lt;0.05; **, p&lt;0.01; ****, p&lt;0.0001; significant difference between saline and drugs).

In females, KOR-mediated alterations were less pronounced. One-way ANOVA showed no drug effects (p=0.4) on signal amplitude (Figure 2D). However, there was an effect on signal width (F(2, 5281) = 19.16; p<0.0001) (Figure 2E), as U50 reduced width (100.0 ± 1.4% vs. 87.7 ± 1.6%; p<0.0001), which was blocked by ATIC (104.8 ± 2.1%; p<0.0001). There was also a significant effect on slope (F(1.209, 7.257) = 7.819; p=0.0225) (Figure 2F), with an increase due to U50 (1.235 ± 0.3967 vs. 1.550 ± 0.4888 %ΔF/F / s; p = 0.0336) that was blocked by ATIC (1.228 ± 0.3919 %ΔF/F / s; p=0.0219). Overall, in males, U50 reduced release and increased uptake of DA, while only increasing uptake in females.

### 3.4. Blocking KORs augments DA signals in naïve animals

After completing U50 trials, the same mice completed trials with ATIC alone and a vehicle control. In males, ATIC increased signal amplitude (100.0 ± 1.1% vs. 117.0 ± 1.8%; p<0.0001) (Figure 3A) and width (100.0 ± 2.0% vs. 124.1 ± 2.7%; p<0.0001) (Figure 3B). A paired t-test revealed no effect on slope (p=0.5530) (Figure 3C). In females, ATIC increased signal amplitude (100.0 ± 0.9% vs. 105.7 ± 1.0%; p<0.001) (Figure 3D) without affecting width (p=0.2809) (Figure 3E). There was a trend towards an increase in slope with ATIC in females (0.9562 ± 0.2592 vs. 1.335 ± 0.4309 %ΔF/F / s; p=0.0847) (Figure 3F). Overall, ATIC increased signal amplitude in both sexes.

**Figure 3:**
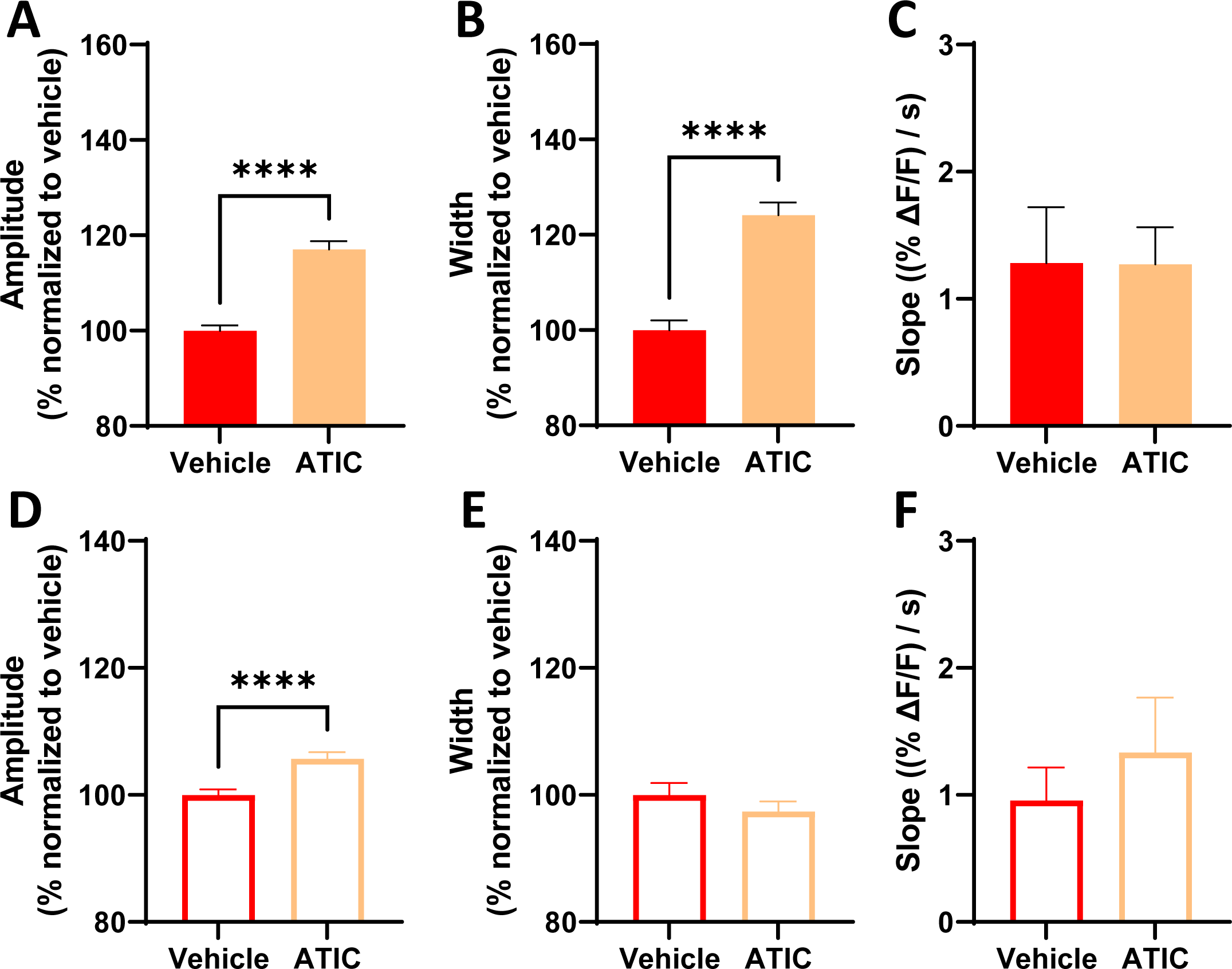
Blocking KORs augments DA signals in naïve animals. Because the mice used herein were naïve to drug, alcohol, and stress insults, it was expected that blocking KORs with ATIC would have no effect on spontaneous signals. In contrast, there was an increase in signal amplitude **(A)** and width **(B)** (normalized to saline control trials) due to ATIC in males. Like the prior raclopride trials (Figure 1), because ATIC caused a proportional increase in amplitude and width, there was no change in slope **(C)** [(ΔF/F (%)) / s]. In females, ATIC also increased signal amplitude **(D)** but there was no change in width **(E)** or in slope **(F).** Thus, ATIC by itself increased DA release without affecting uptake in both sexes. (****, p&lt;0.0001; significant difference between vehicle control and drugs).

### 3.5. KORs modulate the frequency of spontaneous DA signals

The frequency of dLight signals was compared between vehicle and drug trials. In males, one-way ANOVA revealed a treatment effect on signal frequency (F(2,5644) = 14.5; p<0.0001) (Figure 4A). Post-hoc analysis revealed a reduction in frequency due to U50 (0.65 ± 0.011 vs. 0.52 ± 0.024 Hz; p<0.0001), which was blocked by ATIC (0.66 ± 0.017 Hz; p<0.0001). When ATIC was given alone, there was no effect on frequency (p=0.1316) (Figure 4B). Females also showed a significant treatment effect on frequency of signals (F(2, 5757) = 6.500; p<0.01). Post-hoc analysis revealed that U50 reduced frequency (0.62 ± 0.016 vs. 0.53 ± 0.018 Hz; p<0.01) (Figure 4C), which was blocked with ATIC (0.62 ± 0.018 Hz; p<0.01). In contrast to males, females showed an increase in signal frequency when ATIC was given alone (0.57 ± 0.016 vs. 0.69 ± 0.017 Hz; p<0.0001) (Figure 4D). Thus, KOR agonism reduced signal frequency in both sexes, while KOR antagonism promoted greater frequency in females.

**Figure 4:**
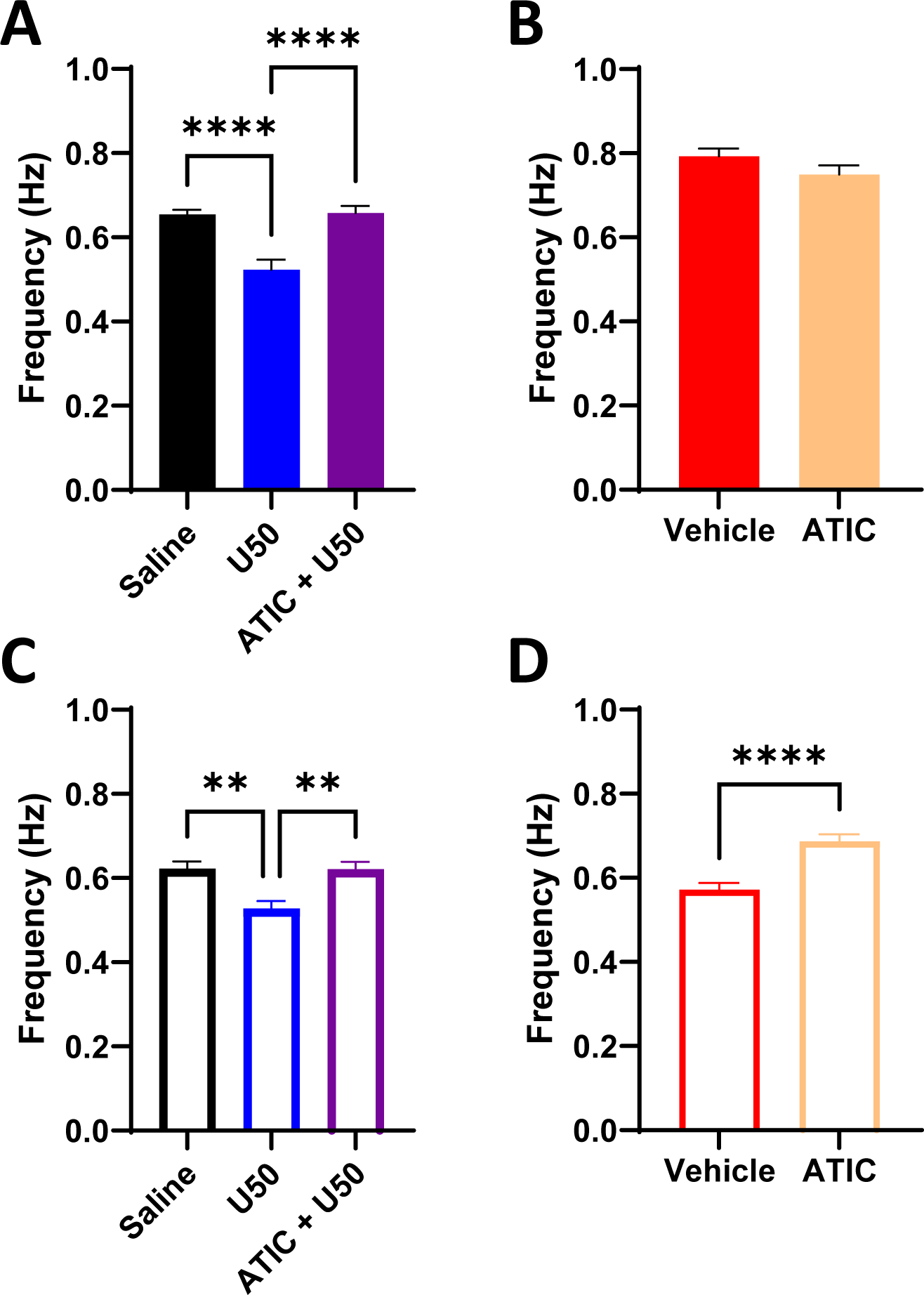
KORs modulate the frequency of DA signals. The peak-to-peak frequency (expressed in Hz) was obtained using pCLAMP, and the resulting values were compared between saline, U50, and ATIC + U50 trials or between ATIC and its vehicle. U50 reduced the frequency of signals in males **(A)** and females **(C).** This effect was blocked by ATIC pretreatment prior to U50. When ATIC was given alone, there was not a significant effect on frequency in males **(B)**, but there was an increase in the frequency of signals due to ATIC in females **(D)**. (**, p&lt;0.01; ****, p&lt;0.0001; significant difference between saline or vehicle control and drugs).

### 3.6. Basal fluorescence intensity levels are reduced by KOR activation

The prior discussions of amplitude and width refer to characteristics of individual spontaneous dLight events. We next took the average fluorescence over an entire recording separately for each excitation wavelength and compared between saline and U50 trials. Representative 15-minute dLight traces of 465 nm-excited fluorescent values are shown for one male (Figure 5A) and one female (Figure 5E), with insets showing respective 405 nm traces. In males, there was a 7.9% reduction of the average 465 nm- excited fluorescence after U50 (159.0 ± 26.8 vs. 146.4 ± 24.4 (arbitrary units); p=0.0403) (Figures 5B, **5C**), but no change in the 405 nm-excited signal (p=0.4736) (Figure 5D). Likewise, in females, there was a 5.4% reduction in 465 nm-excited fluorescence due to U50 (192.2 ± 22.5 vs. 181.1 ± 22.8 (arbitrary units); p=0.0344) (Figures 5F**, 5G**), but no change in the 405 nm signal (p=0.1781) (Figure 5H). This reduction in the 465 (experimental) but not 405 (isosbestic control) channel suggests a KOR-mediated reduction in extracellular DA.

**Figure 5:**
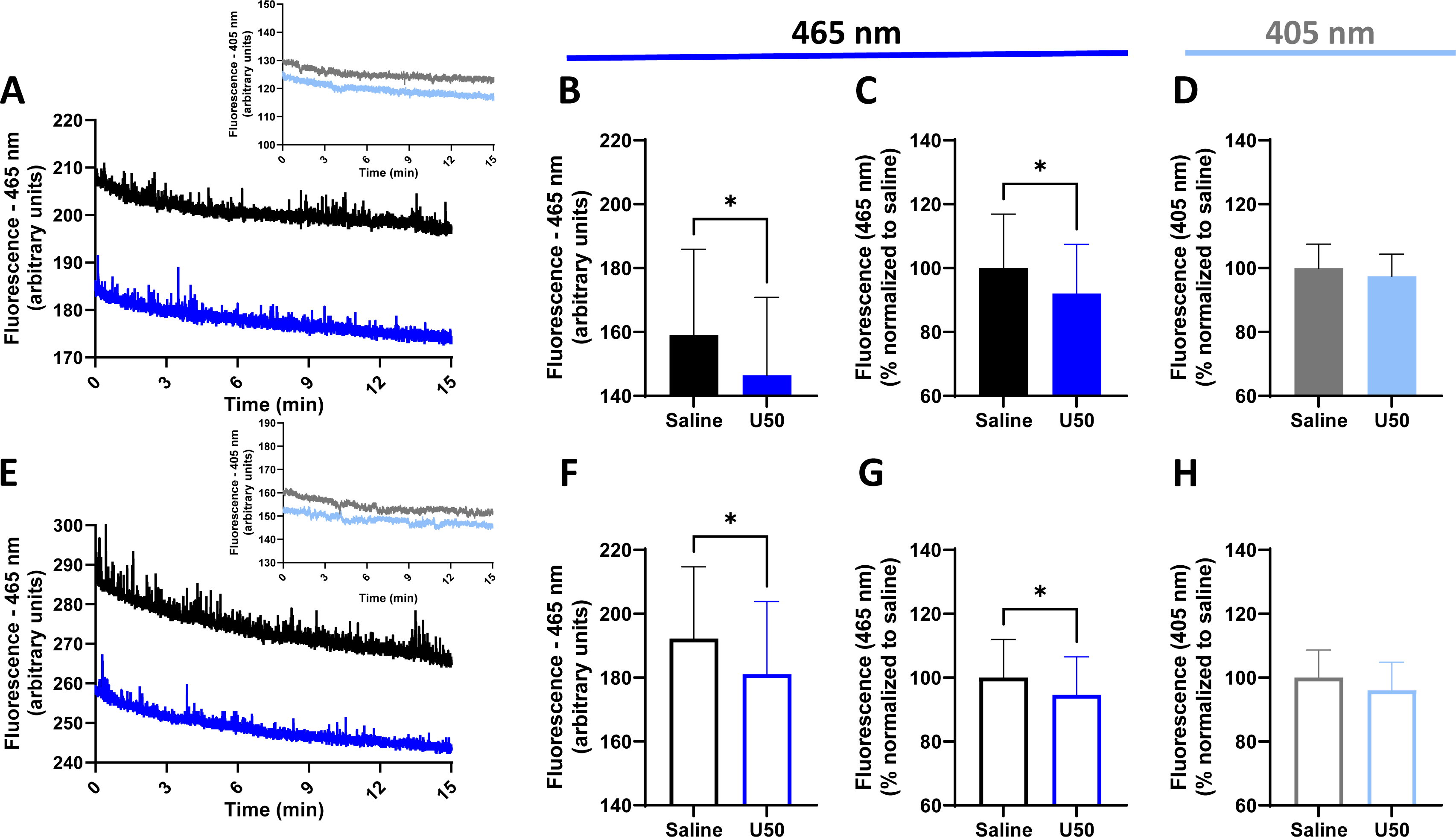
Basal fluorescence intensity levels are reduced by KOR activation. One set each of representative traces showing the actual raw fluorescence values from the 465 nm- excited experimental light between saline and U50 trials is shown for one male **(A)** and one female **(E)**. The insets on each figure show the equivalent raw fluorescence values for the 405 nm-excited control lights from the same respective trials. The average fluorescence over 15-minutes of each of these traces for all animals was taken and compared. U50 reduced the raw 465 nm-excited fluorescence values in males **(B)** and females **(F)** (arbitrary units), which is also shown as a percent normalized to saline for males **(C)** and females **(G)**. In contrast, there was no change in fluorescence from the 405 nm-excited signals in males **(D)** or females **(H)**. Because U50 reduced fluorescence in the 465 nm channel but not the 405 nm channel, this suggests an experimental, pharmacologically-induced reduction in fluorescence separate from any effect of photobleaching. (*, p&lt;0.05; significant difference between saline and drugs).

## 4. Discussion

This study measured endogenous DA in real time in the NAc using dLight and pharmacology to assess KOR-mediated changes in the kinetics of spontaneously arising DA release events. Using drugs with well-established effects on DA, we devised an analysis method to assess amplitude, width, slope, and frequency. Cocaine, a DAT blocker [53], reduced signal amplitude but increased width, leading to a decrease in slope that indicates slower uptake. This aligns with a recent report [35] showing a longer decay constant of spontaneous dLight signals *in vivo* in the dorsolateral striatum due to cocaine. Raclopride, a D2R autoreceptor antagonist, is known to increase NAc DA release [40, 54]. Herein, raclopride increased amplitude and proportionately increased width, suggesting an increase only in DA release. The predictable effects of these compounds on the correlation between amplitude and width (slope) established proof-of-concept that we were assessing release and uptake in spontaneous DA signals.

We reported a reduction in signal amplitude of dLight signals due to U50 in males only. However, both sexes showed a greater reduction in signal width compared to amplitude, resulting in an increased slope, indicative of faster DA uptake. Activation of KORs is known to positively regulate the DAT. Indeed, upregulation of ERK1/2 kinase by KOR activity induced DAT Thr53 phosphorylation [13], increasing DAT trafficking in cells [11, 55] and striatal tissue [13] that promoted faster DA uptake [11, 12], including *in vivo* uptake measured using chronoamperometry [13]. Together with this supporting evidence, our findings suggest that acute KOR activation markedly increases the rate of endogenous DA uptake in freely moving animals. This, however, was a somewhat surprising result, because there is conflicting evidence in the literature on the contribution of increased uptake to KOR-mediated reductions in DA. For example, *ex vivo* experiments in striatal slices [56, 57] and *in vivo* experiments using VTA stimulation [8] showed no difference in DA uptake due to KOR activation. Other studies show negative effects on DA uptake. The KOR agonist nalmefene reduced DA uptake, an effect which was enhanced following chronic ethanol exposure [58]. Further, although acute exposure had no effect, after repeatedly injecting U69,593, the DA extraction fraction in microdialysis samples, an indirect measure of DA uptake, was reduced [14]. Because long-term KOR activation (e.g., via chronic ethanol exposure or repeated agonist injections) has been shown to increase dynorphin expression in the NAc [59] and decrease extracellular levels of DA [60], it is possible that compensatory mechanisms drive slower uptake to restore tonic DA. For instance, low tonic DA would diminish stimulation of presynaptic D2R autoreceptors, which are known to facilitate DAT activity [61], thereby reducing uptake. Further, repeated KOR activation reduced striatal D2R expression [62] and attenuated D2R autoreceptor-mediated reductions in DA release [63, 64]. Thus, effects of KOR activation depend both on prior manipulations of the KOR and compensatory changes to D2R signaling. Our data support that acute, rather than repeated, KOR activation promotes the rate of DA uptake in the NAc *in vivo*.

All three major effects of KOR activation reported herein - reduced release (males), increased uptake (both sexes), reduced frequency (both sexes) - can contribute to a state of hypodopaminergia [65], where tonic extracellular DA concentrations are reduced. The literature is consistent in reporting that KOR activation reduces tonic DA. Microdialysis studies show that KOR agonists, when injected systemically [2, 66] or into the NAc [1, 67], reduce extracellular DA concentrations. In the present study, the hypothesis that background fluorescence levels from dLight sensors reflect “basal” extracellular levels of DA in the brain that can be reduced by KOR activation was determined to be correct. Systemic U50 injection significantly reduced background fluorescence in mice of both sexes, however, females exhibited a smaller decrease in fluorescence. This provides two metrics (amplitude and background fluorescence) by which females were less sensitive to KOR activation.

There are a couple recent reports of changes in background dLight fluorescence. One showed that amphetamine-mediated increases in striatal extracellular DA concentrations measured by microdialysis were similar to increases in background dLight fluorescence [34]. Prolonged elevations in background fluorescence from dLight were also detected after fentanyl injection [36]. Taken together, the present findings suggest that background changes in dLight fluorescence can be used to estimate pharmacologically-induced changes in tonic extracellular DA concentrations.

In many cases, female rodents are less sensitive to KOR-activating drugs. Females exhibited lower KOR-mediated analgesic and antinociceptive efficacy [68, 69], and they were less sensitive to KOR agonist effects on motivated [45] and depressive-like behavior [46]. Further, reductions in DA release due to KOR activation in the NAc *in vivo* [45] and in striatal tissue [70] were smaller in females than males. In contrast, *ex vivo* slice FSCV studies in non-human primates [5, 65] and rodents [4] showed similar KOR-mediated reductions in stimulated DA release in both sexes. In the present study, only males showed reductions in signal amplitude due to U50, supporting that, *in vivo,* males are more sensitive to KOR-mediated inhibition of striatal DA release.

Because we present a novel analysis method in this manuscript, we sought to provide evidence that kinetic variables herein were a valid assessment of DA parameters. One consideration was the potential effect of KOR-mediated changes to neuron firing, which could alter the amplitude and width of signals. It has been shown that firing of NAc-projecting DA neurons was inhibited by KOR agonism in mice [16, 71, 72] but not rats [17, 72]. Because DA release largely depends on action potential generation in DA neurons [73, 74], shorter trains of action potentials [75–77] as a consequence of KOR activation could reduce peak height and make signals look “thinner”. However, the utility of the analysis presented here is the use of slope to measure uptake, which considers changes in both amplitude and width. For example, U50 reduced the amplitude, width, and frequency of signals in males. If effects of KOR activation on DA neuron firing affected amplitude and width, there should be some relationship between them and the frequency of signals. Because the reduction in width was greater than the reduction in amplitude, slope was *increased*, meaning U50 increased uptake while also reducing frequency. A similar increase in slope in females was achieved with U50 through a different relationship between amplitude (no change) and width (decrease), yet frequency was still reduced. On the other hand, ATIC increased signal frequency in females only, though it increased amplitude in both sexes without affecting slope. Thus, these pharmacological manipulations resulted in changes to amplitude, width, and slope that were independent from frequency, supporting that the reduction in frequency of spontaneous signals due to U50 herein is related to KOR-mediated control over DA neuron firing.

One of the most surprising results of the present work was that the KOR antagonist, ATIC, when administered alone, increased the amplitude of dLight signals in both sexes. An antagonist typically should have no effect other than by blocking ligands that would activate the receptor [78]. Therefore, the effect of an antagonist alone often indicates “tone” of the endogenous ligand. It has traditionally been thought that dynorphin tone does not exist at baseline but results from insults such as stress or drugs of abuse [79]. For example, increased dynorphin expression in the NAc has been shown *after* stress exposure [80], and KOR antagonism attenuated depressive behaviors post-stress [80] and during morphine withdrawal [81]. However, in *naïve* C57BL/6J mice, KOR antagonism increased DA release in the NAc [82]. KOR antagonism was also shown to increase DA release without affecting uptake 1 hour after norBNI treatment [83], directly aligning with our timeline and results. This evidence together suggests that dynorphin tone exists *in vivo* in naïve animals.

In conclusion, systemic KOR activation altered the kinetics of real-time endogenous DA signals in a sexually divergent manner. We report an expected KOR-mediated reduction in DA release in males, but increased uptake rates and reductions in frequency of release events in both sexes represent robust and previously underappreciated mechanisms by which KORs regulate DA. The approach offered here provides the groundwork to further explore the complex interactions between KORs and DA. It also provides a means of assessing spontaneous signals in a standardized way, which will be important in determining how this type of signaling might affect behavior.

Different techniques have given disparate views of how KORs modulate DA. By combining measurements of amplitude, width, slope, and frequency of spontaneous dLight signals together in this novel format, a more holistic view of how KOR activation regulates DA in real time may be gleaned. Understanding exactly how KORs modulate DA in an intact animal will help promote the development of effective treatments for pain, depression, and substance use disorders.

## Author Contributions

Conceptualization: SRJ, KMH, CWW; Methodology: CWW, SWC, SRJ, KMH; software: CWW, SWC; Validation: CWW, SWC, SRJ, KMH, CYS; Formal Analysis: CWW; Investigation: CWW, CYS; Resources: SRJ, KMH, SWC; Data Curation: CWW; Writing – Original Draft: CWW; Writing – Review and Editing: SRJ, KMH, CWW, SWC; Visualization: CWW, SRJ, KMH; Supervision: SRJ; Project administration: SRJ, KMH, CWW; Funding acquisition: SRJ, KMH.

---

**Figure S1:**
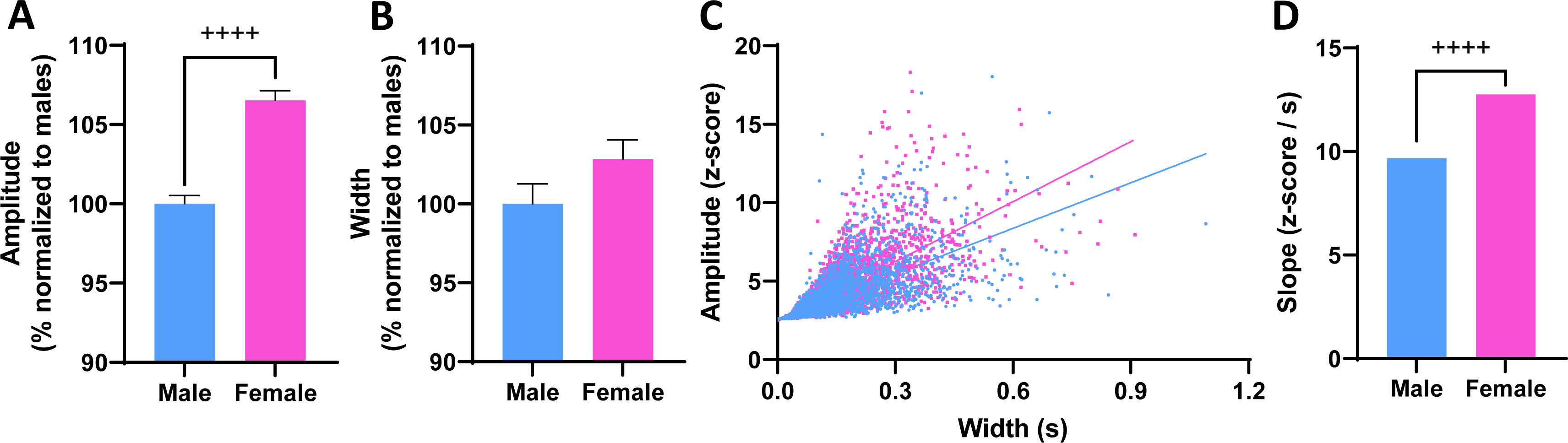
Sex differences in release and uptake. All other figures and analyses are based on a within- animals repeated measures drug design. In this supplemental figure, analyses were conducted using the adaptive iteratively reweighted Penalized Least Squares (airPLS) algorithm. This was done because the strength of raw fluorescence values by light excitation, which affects the amplitude of spontaneous signals, is different between animals. To compare fluorescence values and assess basal uptake metrics between animals, this code was used to normalize ΔF/F values into z-scores for each animal. Using this method, females were shown to have spontaneous signals with greater signal amplitudes (expressed as a percent normalized to males) during saline control trials **(A)**, but signal width did not differ between sexes **(B)**. For every signal detected during saline trials, amplitude values are plotted against width for all males (n=7) and females (n=7) **(C)**, and simple linear regression showed females had a higher slope (z-score / s) than males **(D)**. This suggests that females have faster DA uptake at baseline during control trials. (^++++^, p&lt;0.0001; significant difference between males and females).

