## Supplemental Figure 1 for "Kappa Opioid Receptors Negatively Regulate Real Time Spontaneous Dopamine Signals by Reducing Release and Increasing Uptake"

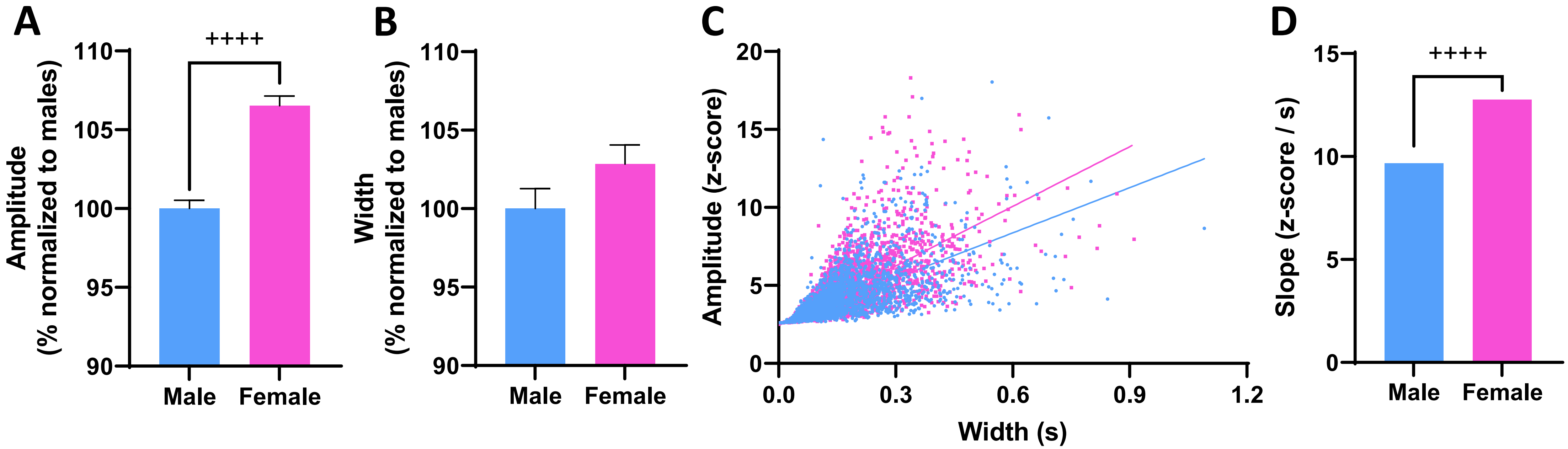


**Figure S1: Sex differences in release and uptake.** All other figures and analyses are based on a within-animals repeated measures drug design. In this supplemental figure, analyses were conducted using the adaptive iteratively reweighted Penalized Least Squares (airPLS) algorithm. This was done because the strength of raw fluorescence values by light excitation, which affects the amplitude of spontaneous signals, is different between animals. To compare fluorescence values and assess basal uptake metrics between animals, this code was used to normalize $\Delta F/F$ values into z-scores for each animal. Using this method, females were shown to have spontaneous signals with greater signal amplitudes (expressed as a percent normalized to males) during saline control trials **(A)**, but signal width did not differ between sexes **(B)**. For every signal detected during saline trials, amplitude values are plotted against width for all males (n=7) and females (n=7) **(C)**, and simple linear regression showed females had a higher slope (z-score / s) than males **(D)**. This suggests that females have faster DA uptake at baseline during control trials. ($^{++++}$, $p<0.0001$; significant difference between males and females).
